# A geniculocortical circuit model predicts functional organization underlying efficient spatial frequency coding in the mouse visual system

**DOI:** 10.64898/2026.09.15.751823

**Authors:** Ahmad Elsayed, Jianhua Cang, C Daniel Meliza

## Abstract

Neural encoding of sensory information becomes more efficient when distinct stimuli evoke uncorrelated activity patterns. Spatial frequency (SF) information becomes decorrelated early in visual processing in a manner that involves both temporal differentiation in the dorsal lateral geniculate nucleus (dLGN) and firing rate differentiation in the primary visual cortex (V1). However, the specific functional circuit configurations that enable this synergy are not fully understood. To address this gap, we developed a geniculocortical subunit model to identify functional circuit configurations capable of producing a decorrelated V1 population response. We found that decorrelation could be explained by the combined activity of two complementary circuit configurations with distinct functional organization. The first circuit was characterized by temporally organized, late-onset dLGN inputs coupled to weak and delayed intracortical inhibition, whereas the second was characterized by less temporally constrained, early-onset dLGN inputs coupled to strong and early intracortical inhibition. Eliminating SF selectivity in upstream dLGN subunits did not affect SF decorrelation in the modeled V1 population response, indicating that decorrelation tolerates weak SF selectivity and is primarily driven by temporal organization. Together, these results suggest that biologically plausible circuit configurations are likely to produce SF decorrelation in the cortex through the temporal organization of feedforward inputs and intracortical inhibition.

**Author Summary:** When visual information enters the early stages of the visual system, it elicits patterns of neuronal activity that encode stimulus features. These codes are more efficient and support better feature discrimination when different stimuli elicit distinct and uncorrelated activity patterns. Population activity patterns depend on how neurons with different functional properties are organized within circuits. We wanted to understand how the functional configuration of circuits connecting the dorsal lateral geniculate nucleus and primary visual cortex enable the decorrelation of spatial frequency-evoked activity patterns. We explored these configurations in a simple computational model that simulates joint geniculate and cortical activity via connected subunits. The model predicted two complementary circuit configurations that recreated the most important underlying dynamics for differentiating SF-evoked activity. These configurations were primarily constrained by the temporal characteristics of geniculate responses and cortical inhibition rather than how strongly individual thalamic units responded to specific spatial frequencies. These findings provide testable predictions about the organization of visual circuits and highlight the importance of temporal organization in efficient coding.

## Introduction

As animals view their environment, their brains take in rich and highly complex visual information. How the visual system transforms this information into a neural code that supports reliable detection and discrimination of biologically salient visual objects is a major question in systems neuroscience. Classical theories posit that stimulus identity is encoded in populations of neurons that respond selectively to only a narrow range of stimuli (1–5). Under this framework, decoding involves pooling across a neuronal population and quantifying the degree to which selective neurons have “voted” for a particular stimulus (6). Decoding accuracy improves as additional neurons are added to the population; however, these improvements have an upper limit due to shared noise across units (7). This limitation suggests that selectivity alone provides an incomplete explanation of the population code.

Alternatively, stimulus identities can be encoded by relationships between their evoked population-wide activity patterns (8–10). These patterns can be conceptualized geometrically as vectors within an N-dimensional state space, where dimensions correspond to neurons and coordinates correspond to firing rates, although the population activity may be confined to a lower-dimensional manifold within this space (4,11,12). Under this framework, stimuli are more differentiated and easier to decode when their evoked response vectors are uncorrelated (i.e., orthogonal, 13–16). Reductions in population correlations (decorrelation) coincide with attention, precede increases in discriminative information, and coincide with improved behavioral performance on a delayed match-to-sample task (15,17). This suggests that the visual system is organized to decorrelate important visual feature representations in the population geometry, facilitating downstream decoding during cognitively demanding visual tasks. However, the biological mechanisms that produce decorrelation are not fully understood.

Prior research suggests a link between decorrelation and coarse-to-fine processing (CTF), a widely conserved phenomenon in which low spatial frequency (SF) stimuli elicit earlier responses than high SF stimuli (18–23). Evidence indicates that CTF emerges in the dorsal lateral geniculate nucleus (dLGN) and is inherited in the primary visual cortex (V1) (18,24,25). CTF occurs during naturalistic behavior (26), coincides with decorrelation (23), and coincides with improved visual category identification (27). Alternatively, delayed nonlinear suppression in V1 has been shown to enhance stimulus selectivity (5,28,29) and promote population sparseness (16,30), and classical theories link these transformations to decorrelation (31–34). However, a recent study from our group indicated that neither geniculate nor cortical mechanisms alone could explain decorrelation. Instead, decorrelation appears to emerge when suppression is temporally aligned with CTF, enabling a geniculocortical synergy in which dLGN and V1 segregate SF information by time and firing rate respectively (16).

Although our previous work revealed that geniculocortical synergy is needed for decorrelation, the functional organization of the dLGN to V1 circuit that produces this synergy remains unclear. Specifically, given the diversity of dLGN response channels (35) and inhibitory V1 dynamics (36), what functional configurations of the circuit produce SF decorrelation? Here, we addressed this question using a simple subunit model of the dLGN to V1 circuit. We confirmed that both CTF and intracortical inhibition are necessary for decorrelation using *in silico* ablation experiments. Our results show that decorrelation can be explained through the combined activity of two circuits with different functional configurations: one characterized by a sequential temporal organization of dLGN afferent responses and one characterized by the strength and timing of intracortical inhibition. We also found that decorrelation persists when SF selectivity in dLGN afferents is eliminated, suggesting that temporal dynamics in the circuit can create a robust V1 population code even when upstream stimulus selectivity is weak or nonexistent.

## Results

### SF decorrelation occurs on a low dimensional manifold

We applied dimensionality reduction via principal components analysis (PCA) to previously recorded responses of 259 V1 neurons to gratings of varying SF (16,23). SF decorrelation was captured with the first two principal components (PCs), which accounted for 24.4 ± 2.2 % (mean ± SD) of the total variance (Fig 1A, B). Both components exhibited coarse- to-fine processing and delayed suppression, but there was an earlier onset in the second component that produced a difference in phase. This delay led to a moment when SF responses were split across the baseline in PC1 and both suppressed below the baseline in PC2. Geometrically, this meant that the response vectors for high and low SF became orthogonal within this low-dimensional manifold (Fig 1C, D). The profiles of the components captured defining characteristics of decorrelation and therefore served as targets for modeling.

**Figure 1.**
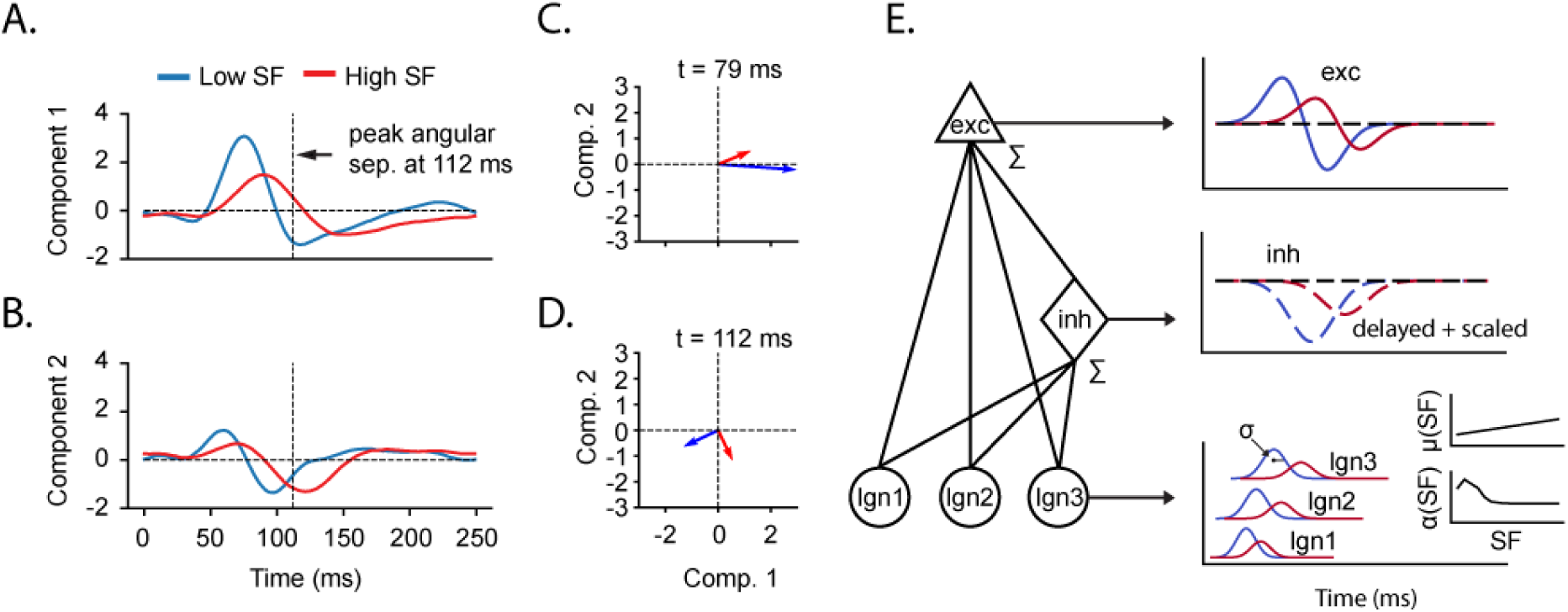
Latent dynamics of V1 SF responses and model architecture. **A.** First principal component (PC1) of V1 responses (n = 259 neurons) to low (0.02 c/d, blue) and high (0.32 c/d, red) SF gratings. Dashed lines indicate the moment when the SF vectors have the highest angular separation (i.e., the late phase). **B.** Same as A, but for PC2. **C.** Response vectors for the lowest (blue) and highest (red) SF in the latent subspace (PC1 and PC2) during the early phase of the response (peak dot product, 79 ms). **D.** Same as C, but for the late phase of the response (peak angle, 112 ms). **E.** Schematic of the model architecture (left) and a visual representation of the output at each stage (right). Subunit identities from bottom to top: dLGN subunits, inhibitory V1 subunit, terminal excitatory V1 subunit. The insets of the bottom panel describe the temporal tuning (*μ*(SF)) and SF amplitude tuning (*α*(*SF*)) functions of the dLGN subunit responses (See methods: Model architecture).

To understand the functional organization that produces SF decorrelation, we constructed a model based on a simple linear combination of geniculocortical subunits with independent tuning and feedforward inhibition in downstream cortical subunits. The circuit was reduced to the minimum hypothesized elements needed to explain decorrelation (Fig 1E). The SF-dependent responses for each dLGN subunit were computed as gaussian functions in a 250 ms temporal domain (Fig 1E, bottom panel). The duration of the response was determined by the bandwidth (σ); the peak response time and peak firing rate were determined by separate linear and gaussian SF-dependent functions (μ(SF), the temporal tuning function; and α(SF), the amplitude tuning function) respectively (Fig 1E, bottom panel insets). Responses from 3 dLGN units were summed by downstream excitatory and inhibitory V1 subunits. The activity of the inhibitory subunit was delayed and scaled (Fig 1E, middle panel) before being applied to the excitatory subunit activity, leading to a biphasic response progressing from enhancement to suppression (Fig 1E, top panel). The biphasic excitatory V1 subunit activity served as the terminal output, which was fit to target responses in subsequent analysis.

### Model optimization finds a stable solution that captures decorrelation

We began by fitting the model to the first two principal components of our V1 recording, with a separate circuit (circuit 1 and circuit 2) for each component (Fig 2A). A multi-start, gradient-based optimization protocol was used and 4000 optimization runs were done in parallel with different random parameter initializations, producing multiple independent converged solutions (see methods: Model optimization). Each solution was used to make a terminal response; the average closely captured the spatiotemporal characteristics of the target (Fig 2B) and 3725/4000 solutions had a loss under 0.15 MSE (Fig 2C). All solutions were included in subsequent analysis.

**Figure 2.**
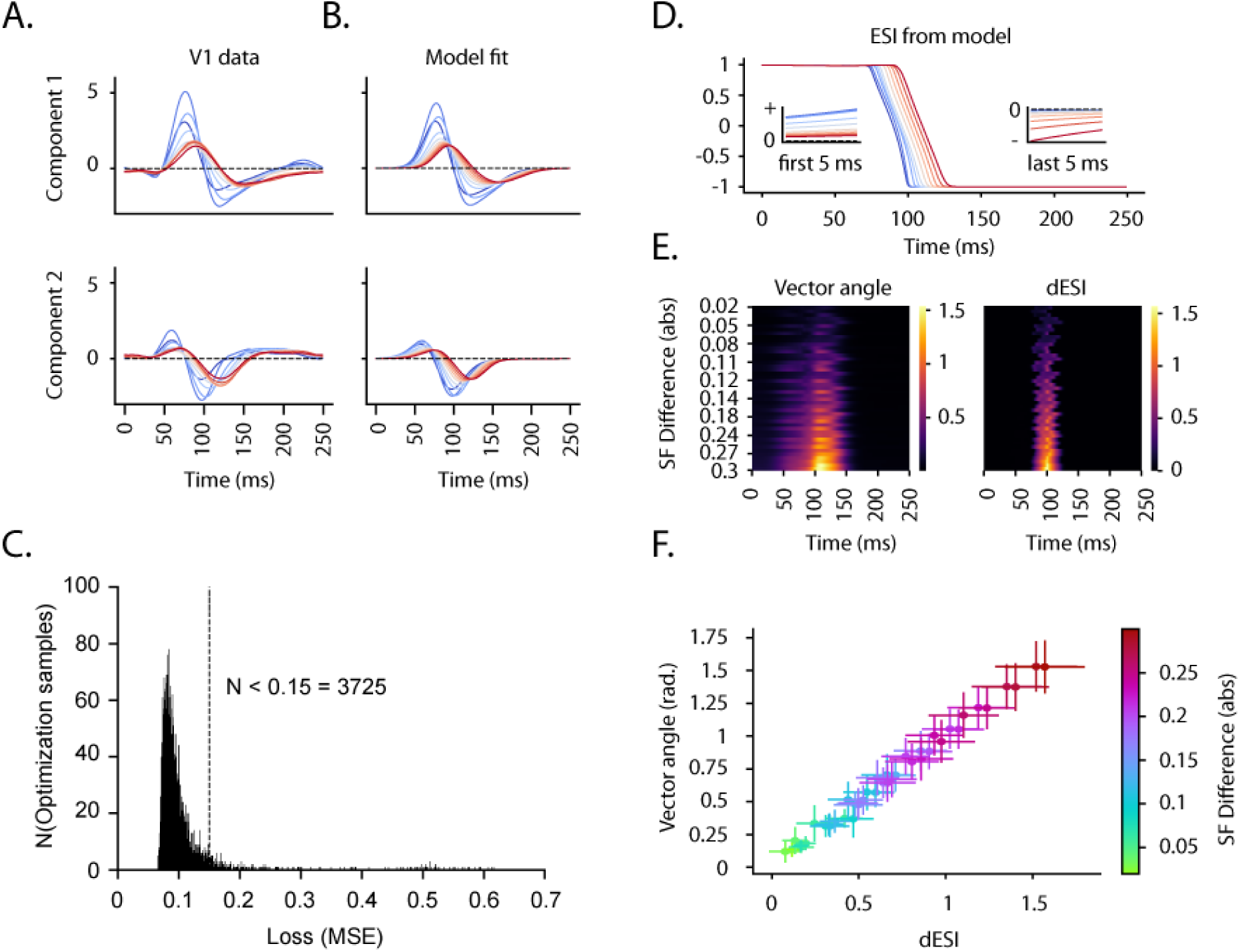
Model optimization and SF decorrelation. **A.** Target PC1 (top) and PC2 (bottom) response profiles to all SF conditions (Blue = lower SF, red = higher SF). **B.** Average of fitted model responses (n = 4000) to the target response profiles shown in A. **C.** Distribution of loss values (MSE) across model solutions after fitting. A significant proportion yielded a loss under 0.15 (93.12 %, X^2^ = 2975.6, p <0.0001). **D.** Enhancement/suppression index (ESI) calculated from model excitatory V1 subunit population response. Color legend is identical to the one in A. Insets: A zoom-in on the first 5 ms (left) of the response showing pure enhancement (ESI = 1), and the last 5 ms (right) of the response showing pure suppression (ESI = -1). This is because there are no baseline fluctuations from noise in the model. **E.** Heatmaps comparing pairwise SF vector angles (left) and pairwise SF-evoked ESI differences (dESI, right). Rows in the heatmap are arranged from top to bottom in order of absolute difference of compared SF conditions in c/d. **F.** Scatterplot comparing SF vector angles (Y-axis) and corresponding dESI (X-axis) at their respective peak times (110 ms and 100 ms respectively; t(3999) = -15.99; p < 0.001, Cohen’s d = - 0.35). At their peaks, dESI and pairwise angles were nearly perfectly correlated (average Pearson R = 0.97). Color indicates absolute difference of compared SF conditions in c/d. Error bars represent standard deviations across optimization runs.

Our previous work revealed a relationship between the dynamic balance of enhancement and suppression and decorrelation in V1. To determine if this relationship was captured by the model, we calculated the enhancement/suppression index (ESI) across the excitatory V1 subunit population for each converged solution. The ESI reflects the ratio between above-baseline and sub-baseline population activity evoked by each SF, where -1 indicates pure suppression, 1 indicates pure enhancement, and 0 indicates equal enhancement and suppression. The average across solutions showed a rapid switch from positive to negative ESI (Fig 2D). During this transition, SFs were temporally differentiated: responses to high SFs remained enhanced for about 20 ms while low SFs became suppressed. We measured the geometric separation between SF vectors over time by their angles (Fig 2E, left). Angles increased with SF difference and peaked at approximately 110 ms. Corresponding ESI differences showed a similar pattern but peaked at approximately 100 ms (Fig 2E, right). At their respective peaks, vector angles and ESI differences were nearly perfectly correlated across model solutions and both increased as the difference between the compared SF pairs increased, consistent with SF decorrelation (Fig 2F).

### Coarse-to-fine processing and intracortical inhibition are necessary and non-substitutable

Our previous study suggested that coarse-to-fine processing and intracortical suppression are both necessary to produce SF decorrelation in V1. We tested this in the model by ablating each hypothesized contributing factor and determining how this affected ESI and the angle between the lowest and highest SF vectors, averaging across converged solutions. The control condition (INH-ON, CTF-ON) showed the characteristic SF-differentiated progression of ESI from enhancement to suppression (Fig 3A, top). The SF angle rose to 45.7 ± 19.1 (mean ± SD) degrees at the early phase (peak dot product) and 87.6 ± 11.6 degrees at the late phase (peak angle) before decaying to 0, showing that the vectors became nearly orthogonal over time (Fig 3A, bottom). When coarse-to-fine processing was ablated (INH-ON, CTF-OFF), ESI progressed from enhancement to suppression, but SF-dependent differentiation was eliminated (Fig 3B, top) and the SF angle peaked at only 15.03 ± 9.9 degrees (Fig 3B, bottom). When intracortical inhibition was ablated (INH-OFF, CTF-ON), ESI remained at 1 throughout the full course of the response and the SF angle rose to a peak of only 24.4 ± 11.6 degrees (Fig 3C). Ablation of both coarse-to-fine processing and suppression (INH-OFF, CTF-OFF) resulted in ESI fixed to 1, and the SF angle remained near 0 throughout the entire response window (Fig 3D). For all experimental conditions, the late phase (peak) angle was lower compared to both the early and late phase angles of the control condition (S1 Supplementary Material).

**Figure 3.**
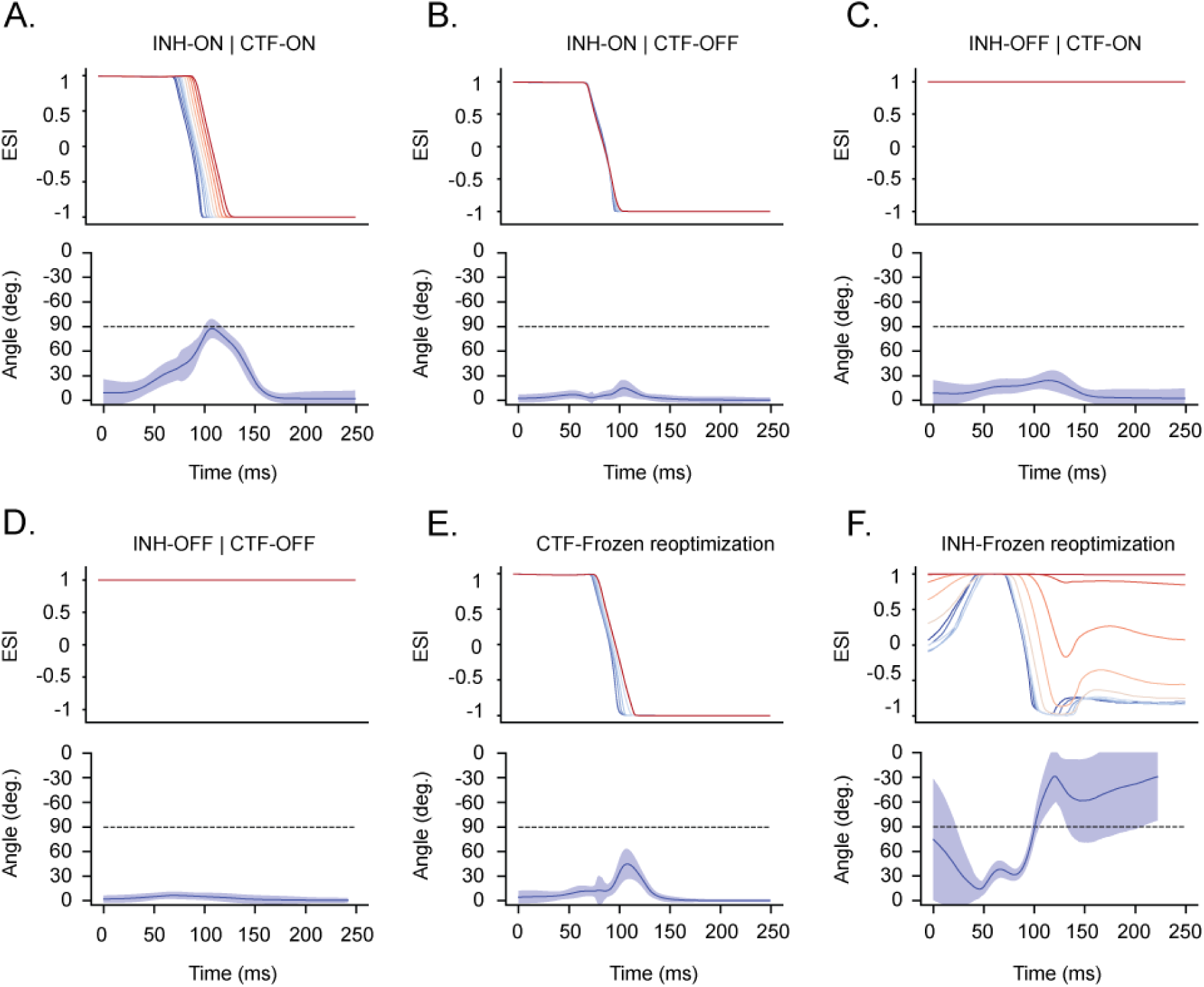
Ablation and fixed optimization experiments. **A.** The baseline condition in which coarse-to-fine processing and intracortical inhibition are both on. Top panel shows ESI as a function of time after stimulus onset (same as Fig. 2D, but insets removed). Bottom panel shows mean (± SD) angle between response vectors for the lowest and highest SF (identical to the bottom row of the left heatmap in 2E). All subsequent panels follow the same format. **B.** Inhibition-on, coarse-to-fine-off ablation condition. **C.** Inhibition-off, coarse-to-fine-on ablation condition. **D.** Inhibition-off, coarse-to-fine-off ablation condition. **E.** Results from the frozen coarse-to-fine processing optimization experiment. **F.** Frozen intracortical inhibition optimization experiment. All statistical comparisons included in (S1 Supplementary Material).

Two additional optimization conditions were tested to determine if coarse-to-fine processing and intracortical inhibition could be replaced by other mechanisms in the model. In the first condition (CTF-Frozen), the frequency-time slope (FTS, model parameter controlling coarse-to-fine strength) was fixed to 0 to determine if alternate configurations of dLGN subunit onset time and response duration can compensate for the loss of coarse-to-fine dynamics. ESI followed a similar progression from enhancement to suppression as in the baseline condition, but the SF-dependent differentiation was diminished (Fig 3E, top). The resulting SF vector angle peaked at 44.87 ± 18.27 degrees, similar to the early phase of the control (Fig 3E, bottom, S1 Supplementary Material). In the second condition (INH-Frozen), inhibition weight was fixed to 0 and dLGN responses were allowed to be negative, corresponding to non-preferred phases evoking sub-baseline firing rates through withdrawal of excitation (37,38). In this condition, suppression was asymmetric across SF conditions, preferring mainly low SFs. As a result, ESI remained fixed to 1 for the highest SF condition and most high SF conditions were only partially suppressed throughout the entire response window (Fig 3F, top). The resulting SF vectors were colinear but had opposite magnitudes (Fig 3F, bottom).

### dLGN temporal organization and intracortical inhibition are asymmetric across circuits

We next sought to compare how subunits with different functional properties were organized within the two circuits (circuit 1 fit to PC1; circuit 2 fit to PC2), starting with the V1 layer. Inhibitory weights in circuit 1 were smaller than in circuit 2 (Fig 4A, left), whereas inhibitory delays were later in circuit 1 than in circuit 2 (Fig 4A, right). These asymmetric configurations reflected the respective geometric profiles of the PCs; PC1 required a relatively weaker, delayed inhibitory drive to split low and high SF responses across the baseline, whereas PC2 required a strong, early inhibitory drive to push all SF responses below the baseline.

**Figure 4.**
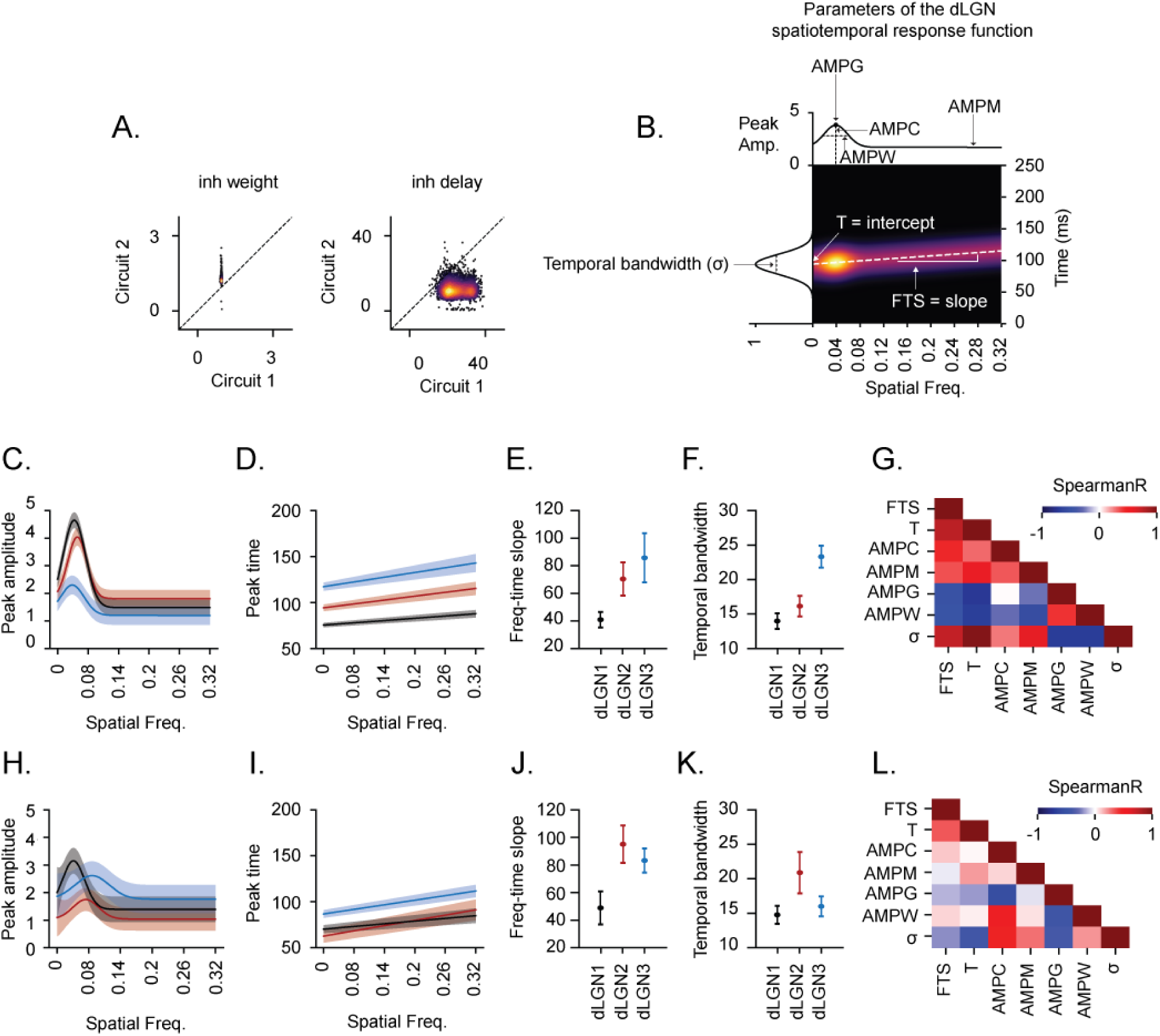
Model circuit configurations. **A.** Distribution of estimates for inhibition weight (left) and inhibition delay (right) in circuit 1 and circuit 2, across 4000 optimization runs with varying initial conditions. **B.** Illustration of how the spatiotemporal response of a dLGN unit depends on the amplitude (top) and temporal (white dashed line) tuning functions. Left: The gaussian temporal envelope of the dLGN response. Parameter abbreviations: FTS = Frequency-time slope; T = Temporal intercept; AMPC = Amplitude center; AMPG = Amplitude gain; AMPW = Amplitude width; *σ* = Temporal bandwidth. **C.** Amplitude tuning functions (used to compute peak SF-evoked firing rate) for the circuit 1 dLGN afferents. Average (solid line) and SD (shaded area) across 4000 optimization runs is shown. Each color corresponds to a different dLGN afferent (Black: dLGN1, Crimson: dLGN2, Blue: dLGN3). **D.** Temporal tuning functions for circuit 1 afferents. **E.** Frequency-time slopes (used to compute coarse-to-fine shifts) for circuit 1 afferents. **F.** Temporal bandwidths (used to compute response duration) for circuit 1 afferents. **G.** SAMs for the circuit 1 afferents Exact Spearman R values and statistical comparisons included in (S2 Supplementary Material). **H-L:** Same as C–G but for circuit 2 afferents.

We next analyzed the tuning properties of the dLGN afferents (Fig 4B). The values of different dLGN parameters can covary across the 3 circuit subunits in many ways, resulting in different permutations. For example, response latencies may increase or decrease with amplitude gain, or may be largely unrelated. Therefore, averaging across unconstrained model solutions obscures meaningful co-ordering structure. To obtain identifiable solutions for exploring the dLGN parameter space, we did a separate optimization (4000 parallel runs) where solutions were constrained to the permutation that yielded the lowest average loss (see Methods: Cross-parameter co-ordering permutation search, S2 Supplementary material).

The dLGN afferents in circuit 1 exhibited a functional temporal sequence. The amplitude tuning functions were all narrowband and had similar preferred SF of 0.05 – 0.06 c/d (Fig 4C) but differed in peak response amplitude. The temporal properties negatively covaried with the peak amplitude: the subunit with the highest amplitude had the earliest peak time (Fig 4D), weakest coarse-to-fine dynamics (Fig 4E), shortest duration (Fig 4F) and vice versa. To confirm this result was not an artefact of the constrained fit, we calculated Spearman rank-order correlations between each pair of parameters for the 120 lowest loss unconstrained solutions (see methods: Cross-parameter co-ordering permutation search). This yields 7x7 Spearman autocorrelation matrices (SAMs) where each cell represents the covariance of two parameters across the 3 dLGN afferents of a circuit. The average SAM revealed similar circuit 1 configurations across the 120 best unconstrained solutions (Fig 4G).

In contrast, the circuit 2 afferents exhibited less clear functional organization, suggesting the strong and early intracortical inhibition may be more important than the upstream input pattern for producing the necessary terminal response profile. One amplitude tuning function was low-pass and narrow band while the other two were slightly more high-pass and had slightly broader tuning (Fig 4H). Furthermore, the circuit 2 afferents had earlier latencies than those in circuit 1, reflecting the temporal profile of the PC2 target (Fig 4I). Both the amplitude tuning functions and temporal tuning functions of the circuit 2 afferents were less differentiated compared to circuit 1 (Fig 4I-L).

### Decorrelation tolerates non-SF tuned dLGN inputs

Previous literature suggests that weak stimulus selectivity in single neurons does not prevent decorrelation on a population level (17,39,40). This raises the possibility that decorrelation depends mainly on the temporal organization of dLGN inputs to V1, without requiring SF-selective responses. To test this, the model was optimized with flat amplitude tuning curves in the dLGN subunits (i.e., peak response rates were identical across SF conditions), eliminating SF selectivity. All other parameters were allowed to freely converge during optimization.

Eliminating SF selectivity in the dLGN subunits had little effect on the temporal characteristics of the terminal V1 response profiles (Fig 5A). In these, the amplitude across SF retained its low pass characteristics due to upstream coarse-to-fine processing reducing the temporal overlap of upstream high SF responses. The asymmetry of inhibition between circuit 1 and circuit 2 was still present, and the temporal parameters of the dLGN afferents maintained their sequential organization in circuit 1 (S3 Supplementary Material). The ESI dynamics closely resembled those from the original optimization (Fig 5B, top) and the angular separation between the lowest and highest SF vector was largely preserved, peaking at 84.75 ± 13.66 degrees (Fig 5B, bottom, S2 Supplementary Material).

**Figure 5.**
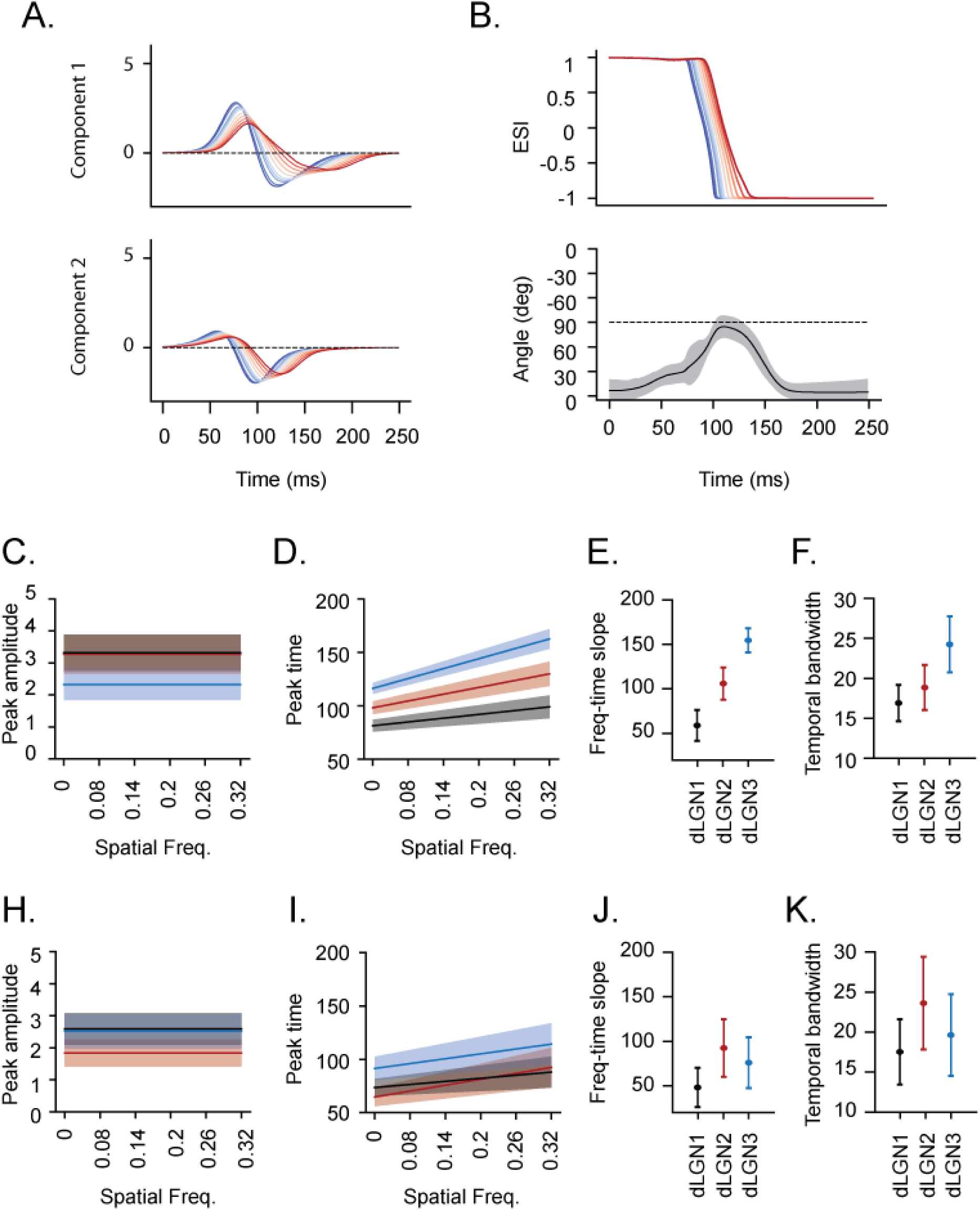
Model results after eliminating SF selectivity in dLGN afferents. **A.** Response profiles from circuit 1 (top) and circuit 2 (bottom) with flat SF-amplitude tuning in upstream afferents. **B.** ESI (top) and SF vector angle between lowest and highest SF (bottom) with flat SF-amplitude tuning in upstream afferents. **C-K.** Amplitude tuning functions, temporal tuning functions, frequency-time slopes, and temporal bandwidths for dLGN afferents of circuit 1 and circuit 2. These are similar to Fig. 4C-F, H-K but for the flat amplitude tuning function experiment.

To assess whether parameter values adjusted to compensate for the elimination of SF selectivity, the tuning functions from this optimization condition were analyzed after applying the same identifiability constraint from the previous section. Compared to the original optimization, the peak amplitudes across circuit 1 afferents decreased, while those from circuit 2 did not change (Fig 5C, H). The response onset times were similar in both circuits (Fig 5D, I).

Coarse-to-fine strength increased in all circuit 1 afferents but only increased in a single circuit 2 afferent (Fig 5E, J). Finally, response durations were marginally higher in both circuit 1 and circuit 2 afferents (Fig 5 F, K). Together these results indicate that decorrelation survives the elimination of SF selectivity in the model, and that the model exploits the temporal organization of the dLGN afferents slightly more when SF selectivity is eliminated.

### Out-of-sample prediction of in vivo dLGN properties

Up to this point, the parameters of the model have been inferred by constraining the output to match the first two PCs of our recorded population. To validate these inferences, particularly the parameters related to dLGN tuning, we fit the model separately to each of the 259 V1 units in the dataset and compared the distribution of dLGN parameters to direct measures from previous *in vivo* recordings from 237 dLGN units (16). Note that whereas the model predicted onset time and response duration from the mean and standard deviation of the gaussian response envelope, the much noisier *in vivo* data requires using noise-threshold-crossing event times. We estimated these threshold-crossing times using the temporal bandwidth (standard deviation) and *in vivo* response signal-to-noise ratio (see methods: Fitting the model to V1 single units).

The model achieved good fits to the individual V1 single unit responses, indicating that it can capture a broad range of V1 response profiles beyond the latent dynamics in the PCs (Fig 6A, B). For the dLGN parameters, there was good agreement between the model and the data for frequency-time slopes and onset times (Fig 6C, D). The model predictions for response duration were systematically lower than *in vivo* data, with an average discrepancy of 2.4 ms (Fig 6D).

**Figure 6.**
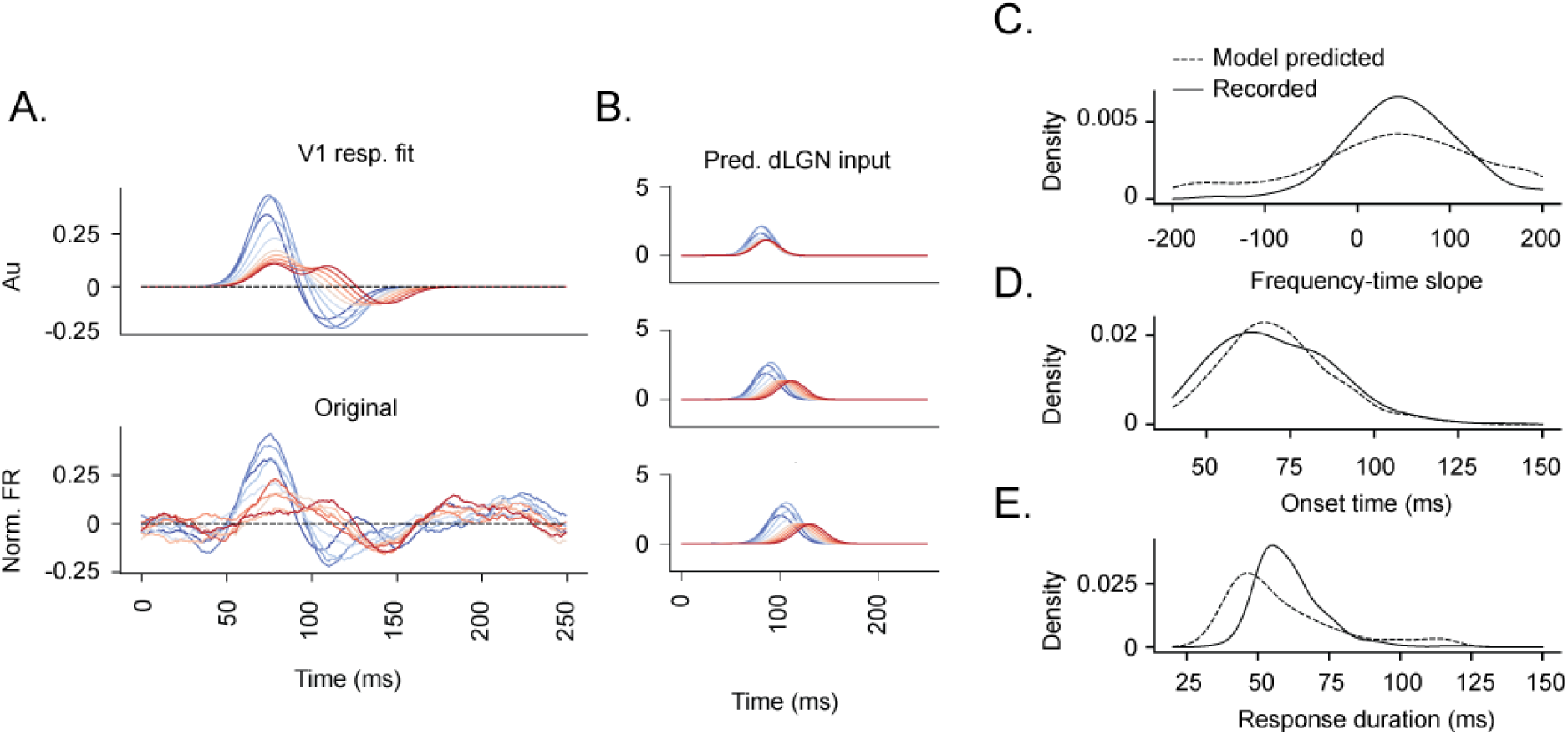
Model dLGN predictions compared to in vivo estimates. **A.** Example of model fit to a single unit V1 response. Top left: model predicted V1 response. Bottom left: Original V1 single unit response. **B.** Each model predicted dLGN response used to make the predicted V1 response in **A**, all fits available to view in the GitHub repository linked in the Data Availability Statement. **C.** Comparison of model predicted frequency-time slopes and in vivo estimates (Mann-Whitney U = 85804, p = 0.11, Cohen’s d = -0.18). **D.** Comparison of model predicted onset times and in vivo estimates (Mann-Whitney U = 86499, p = 0.16, Cohen’s d = -0.19). **E.** Comparison of model predicted response durations and in vivo estimates (Mann-Whitney U = 69552, p < 0.001, Cohen’s d = -0.13).

This divergence could reflect the model’s temporally symmetric response envelope, whereas true responses tend to be asymmetric with longer decays. Overall, these results validated that the model could make biologically meaningful predictions of upstream dLGN inputs to V1.

## Discussion

Our study sought to understand how the geniculocortical circuit produces decorrelated representations of spatial frequency. We hypothesized that a simple model with linear summation and feedforward inhibition would be sufficient to produce SF decorrelation and to infer the functional circuit configurations that facilitate it. To test this hypothesis, we fit the model to the first two principal components of the V1 population response, a sufficient dimensionality to observe SF decorrelation (Fig. 1). Despite strong constraints on the connectivity and functional tuning, the model accurately captured the characteristics of the target PCs and produced a decorrelated population response (Fig. 2). We then showed that coarse-to-fine processing and inhibition as implemented in the model are both necessary for decorrelation and non-substitutable (Fig. 3).

The model parameters consistently converged on configurations in which circuit 1 and circuit 2 had asymmetric dLGN and inhibitory properties. The circuit 1 afferents were temporally lagged, and inhibition was weak and delayed while the circuit 2 afferents were less lagged with strong and early inhibition. In circuit 1, afferent responses were sequentially ordered such that earlier units had higher peak firing rates, weaker coarse-to-fine dynamics, and shorter responses, while later units exhibited the opposite pattern. In circuit 2, afferents had more overlapping functional properties (Fig 4). When we flattened the SF-amplitude tuning curves of the dLGN subunits, we saw almost no effect on decorrelation (Fig. 5), indicating that decorrelation in the model depends more on the temporal organization of dLGN inputs rather than their selectivity for specific SF conditions. The model predictions were largely consistent with *in vivo* dLGN estimates from our previous study (Fig. 6).

Together, these results raise important implications and predictions about the neural code in the early visual system and the biological material that supports it. The population activity underlying SF decorrelation was confined to two latent dimensions, suggesting a distributed population code. These latent dimensions were associated with circuits resembling known dLGN biological channels that facilitate temporal organization. The model also suggests that feedforward inhibition in V1 is sufficient to transform temporally organized upstream inputs into a decorrelated SF representation, but does not rule out the possibility of recurrent connections supporting long-timescale representation. Finally, decorrelation tolerated the elimination of SF selectivity in dLGN afferents, suggesting that decorrelation and stimulus selectivity serve distinct but complementary roles in visual processing.

### Low latent dimensionality suggests a decorrelated & distributed population code

Activity in neural populations, though nominally high-dimensional, is often confined to a low-dimensional manifold that balances robustness and discriminability (4,8,41,42). Consistent with this, SF decorrelation was fully captured by the first two principal components in our V1 data (Fig 1A-D). These PCs capture a high degree of shared variability in the population, indicating that SF information is widely distributed across neurons. Consistent with this, our previous study showed that SF decorrelation does not coincide with the sparsest moment of the V1 population response (16). These results suggest that decorrelation and sparseness are not causally linked as previously hypothesized (34,43). One possibility is that decorrelation is a necessary (but insufficient) condition for promoting sparsity and facilitating dimensional expansion when input stimuli are also high dimensional (44–46). A population code that is decorrelated while sparsity remains constrained is likely advantageous in terms of efficiency and reliability. Decorrelation can improve efficiency by geometrically separating stimuli on the neural manifold. Meanwhile, distributed activity can improve reliability by spreading information across neurons (47). Our results also raise the possibility that the latent dimensions reflect functional biological channels rather than mathematical abstractions, which we discuss in depth in following sections.

### Temporal organization across and within dLGN units sets the stage for SF decorrelation

The two circuit configurations identified by the model had dLGN afferents which were primarily characterized by their temporal organization. These resemble *in vivo* lagged and nonlagged dLGN cells respectively, which exhibit significant response lag differences and produce low dimensional latent dynamics (24,35,48–51). Lagged dLGN cells receive early hyperpolarizing GABAergic inputs from the retina that lead to delayed and broadly varying response lags (37,38), consistent with the afferents of circuit 1 (Fig 4D). In contrast, nonlagged cells exhibit early responses with narrow variance (38), consistent with the afferents of circuit 2 (Fig 4I). Furthermore, retinal inputs to lagged cells are slow and low-amplitude (38); this may propagate to inhibitory V1 cells and lead to weaker activation, providing a plausible explanation for the relatively weak and delayed intracortical inhibition in circuit 1 (Fig 4A). Conversely, the model predicts early and strong inhibition in circuit 2 (Fig 4A), suggesting strong and fast excitation from the retina. Experiments manipulating lagged and nonlagged dLGN channels via targeted blockade of GABAergic retinogeniculate inputs could test these predictions directly.

The circuit 1 afferents exhibited a temporal sequence strongly associated with their spatiotemporal response characteristics. Specifically, early subunits had higher peak response amplitudes, weaker coarse-to-fine shifts, and shorter response durations, while later subunits exhibited the opposite pattern (Fig 4C-G). Response onset times of *in vivo* dLGN cells positively covary with receptive field (RF) size (52); furthermore, coarse-to-fine dynamics are associated with reductions in RF size, with stronger shifts linked to greater reductions (19). This raises the possibility that dLGN onset times covary with coarse-to-fine dynamics *in vivo*, consistent with our model’s prediction. Our findings therefore suggest that lagged cell inputs to V1 follow a temporal sequence, and that a unit’s position in the sequence entails specific spatiotemporal response characteristics.

### Constrained feedforward inhibition separates SF representations in V1

The simplified feedforward construction of the model circuit reflects a known biological feedforward inhibition motif: dLGN makes direct contact onto V1 PV interneurons, which in turn synapse onto principal excitatory neurons (53). Our model shows that inhibitory activity in this motif should be highly constrained to segregate stimulus feature representations (Fig 4A); consistent with this, optogenetic activation of PV neurons with precise timing and laser intensity leads to improved behavioral performance in an orientation discrimination task in mice (54). Our model likely captures a feedforward structure embedded in a larger inhibition-stabilized network (ISN), where feedback inhibition and excitatory recurrence interact on longer timescales (55). In V1, these interactions facilitate surround suppression, contrast gain control, and contextual modulation (36,56). Throughout the wider cortex, ISN activity generates persistent attractor dynamics that facilitate memory and retrieval, suggesting that similar ISN activity in V1 plays a role in forming persistent representations of stimuli (55).

### Decorrelation and feature selectivity may serve distinct but complementary roles

Our results revealed that decorrelation can occur when the amplitude of dLGN responses does not depend on SF (Fig 5), suggesting that decorrelation and stimulus selectivity are separate but complementary processes. One possibility is that selectivity contributes less to geometric separation and more to stabilizing feature representations against variability and noise. Separate studies have shown that shared representational noise decreases with dynamic sharpening of stimulus selectivity (1,2,4) and during attentive viewing (17,57,58). These transformations coincide with improved behavioral performance during a stimulus discrimination task (59), suggesting they underlie improved perceptual fidelity for salient visual features. We hypothesize that decorrelation drives the initial mapping of feature representations in the population state space, while stimulus selectivity promotes their stability. Under this hypothesis, physiological attenuation of stimulus selectivity should reduce representational fidelity while preserving geometric separation. In a perceptual discrimination task, this would likely lead to performance decreasing but saturating above chance.

### Model limitations

The coarse-to-fine dynamics of dLGN subunits in our model are inspired by a classical dynamic center-surround spatiotemporal receptive field (STRF) model (19,24) which predicts a linearly increasing onset time with increasing SF value. Future modeling work is required to test other STRF models of dLGN (60). Furthermore, a broad consensus is missing regarding the number of dLGN cells that connect to a single V1 cell. Classical studies initially proposed estimates between 1 - 10 dLGN neurons per V1 neuron (61,62), however subsequent studies expanded the estimate to 30 - 125 (49,63). More recent work has also suggested that geniculocortical convergence is functionally transient and dynamic over time (64). We therefore took a parsimonious approach, using the minimum number of dLGN subunits needed to meaningfully analyze co-ordering relationships across dLGN parameter classes. Each subunit may therefore reflect the characteristics of a separate population of similarly tuned dLGN cells.

Other abstractions were made to keep the model parsimonious. The model is purely feedforward. Omitting recurrent dynamics may obscure long-timescale response characteristics; however, the scope of this study was mainly concerned with short-timescale dynamics. The model is also agnostic of interneuron identity; inhibition in the model is a subtractive, scaled and delayed copy of the same summed dLGN input terminating onto the excitatory subunit. Divisive inhibition may affect how multiple SF response profiles are integrated; therefore, future work examining natural image responses should incorporate all forms of inhibition known to exist in cortex. Furthermore, only two copies of the circuit are optimized to facilitate a low-dimensional analysis. Although these dimensions captured the SF dynamics of interest for gratings, which have narrowband SF spectra, the broader distribution of SFs in natural images may require analysis in higher dimensional or nonlinear manifolds (8). The dLGN response profile is treated as a temporally symmetric gaussian envelope whereas dLGN responses in a biological system are not perfectly symmetric; this may have led to the underestimation of response durations in this study (Fig. 6F).

We used a multi-start gradient descent-based optimization procedure to assess parameter variability across solutions instead of using a Bayesian approach, which would introduce additional computational complexity. Although this gradient-based method does not provide a formal estimate of the posterior uncertainty, it is useful for revealing non-identifiability in the parameter space (65). This was crucial for evaluating the quality of co-ordering permutations in the dLGN afferents which produced distinct non-equivalent minima in the loss landscape. We therefore constrained the model to the co-ordering permutation yielding the lowest average loss (see methods: Cross-parameter co-ordering permutation search). This constraint eliminated permutation-related variability when assessing dLGN parameter variability across model solutions.

### Future directions

Our results identify temporal organization of activity as an organizing principle behind decorrelation. Specifically, decorrelation relies on both the timing and configuration of dLGN inputs as well as the timing and strength of intracortical inhibition. Temporal organization provides a coding dimension beyond firing rate that scales to high-dimensional populations. When temporal offsets are combined with precisely timed firing rate modulation mediated by inhibition, overlapping feature representations can be separated in the population state space. These results highlight the importance of the geniculocortical mechanisms that facilitate temporal organization and identify important biological targets for future research such as lagged vs. nonlagged channels and feedforward vs. recurrent inhibition. In this study, we constructed and tested the model based on responses to sinusoidal gratings, which are dominated by a single SF. However, natural images contain a spectrum of SFs, which may or may not be processed independently. One possibility is that each SF response is weighted by its corresponding spectral power (41). Alternatively, the summed response might be normalized by the cumulative spectral power, a form of variance normalization that can highlight high SF bands which are likely to carry more perceptually relevant information (67). Future work with high-density recording of population responses to natural images and sinusoidal gratings can address this question directly.

## Materials and Methods

### Dimensionality reduction and response geometry analysis

This analysis was done using previously recorded V1 data (16,23). The data consisted of responses from 259 units to sinusoidal gratings varying in orientation (in degrees: 0, 45, 90, 135), phase (in degrees: 0, 90, 180, 270), and SF (in cycles per degree: 0.02, 0.04, 0.08, 0.1, 0.12, 0.16, 0.2, 0.24, 0.28, 0.32). For each joint orientation and phase condition (N = 16), the data are an array of 10 SF conditions x 250 time lags x 259 neurons. For each of these conditions, we reshaped the array to combine the SF and time dimensions (2500 x 259) and applied PCA to extract principal components (PCs) in the neuron space. We averaged the transformed response profiles across the orientation and phase conditions and reshaped the transformed data back to its original shape (10 x 250 x 259). The first two PCs (cumulative variance explained = 24.4 ± 2.2 %, mean ± *SD*) captured the defining characteristics of decorrelation and were used as modeling targets (10 x 250 x 2) (Fig 1A - C).

### Model architecture

The model is a geniculocortical feedforward inhibition circuit with 3 dLGN subunits feeding into an inhibitory and excitatory V1 subunit (Fig 1E). Each dLGN subunit produces a time-varying response that depends on the SF of the input stimulus (in cycles per degree, c/d). Each of the V1 subunits sums upstream dLGN subunit responses; the excitatory V1 subunit also sums the output of the inhibitory subunit, scaled by an inhibitory weight (INH_W_) and delayed by an inhibitory delay (INH_D_). The excitatory V1 subunit produces the terminal output of the circuit (Fig 1E, left).

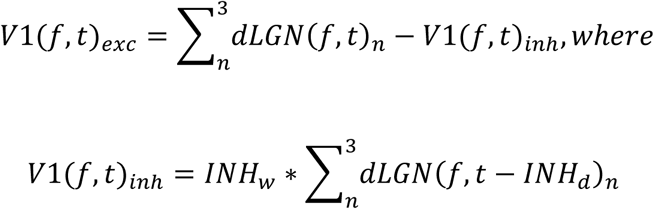

The response of each dLGN subunit is modeled using a gaussian function with a temporal center and amplitude that depend on SF and with temporal bandwidth *σ*.

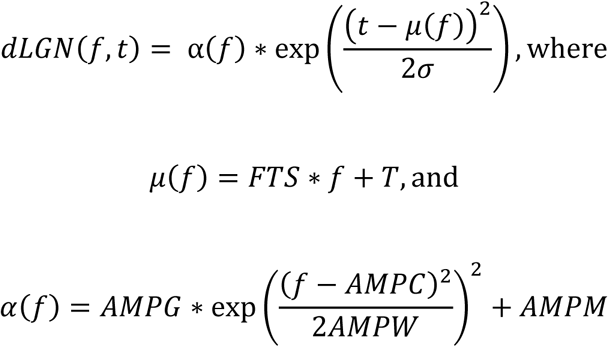

The temporal tuning function *μ*(*f*), which determines the peak response time of the subunit, is a linear function of SF, with the frequency-time intercept (T) controlling the baseline peak response timing and the frequency-time slope (FTS) controlling the temporal shift as SF values increase (i.e., the parameter controlling coarse-to-fine dynamics). The amplitude tuning function *α*(*f*) defines the SF-depedent peak response amplitude, with the amplitude midline (AMPM) controlling the baseline amplitude across the full tuning curve, the amplitude center (AMPC) controlling the preferred SF, the amplitude gain (AMPG) controlling the scaling factor of the preferred SF, and amplitude bandwidth (AMPW) controlling the width of the tuning curve.

Each dLGN response is defined by 7 parameters; the responses across all 3 dLGN subunits are computed from a 7 x 3 matrix in the forward pass. Furthermore, each circuit has a single inhibitory subunit whose dynamics are determined by 2 parameters. Therefore, each circuit is defined by 23 parameters; since two instances of the circuit are initialized and fit in parallel to each of the target V1 PCs, 46 parameters in total are optimized during each fit.

### Model optimization

Model parameters were optimized with a multi-start gradient descent approach (using the ADAM algorithm), with multiple random starting parameters sets initialized and optimized in parallel to minimize the loss function. The loss function was the mean squared error of the model’s terminal output with respect to the target responses, averaged across targets. TensorFlow (version 2.10.1) was used to compute the gradients during the forward pass, and a set of custom Python (version 3.9.23) classes was made to control the full optimization loop. The initial starting parameters were all drawn from the same normal distribution with a mean of 0 and standard deviation of 1 (normalized parameters). During the forward pass, a bounded sigmoid transformation was applied to the normalized parameters to transform them to a biologically plausible range (INH_w_: 0 - 3 Au; INH_d_: 0 – 40 ms; T: 40 – 125 ms; FTS: 0 – 175 ms; *σ*: 10 – 40 Au; AMPM: 0 – 3 Au; AMPC: 0.01 – 0.1 c/d; ampg: 0.1 – 4 Au; AMPW: 0.01 – 0.08 Au). We did 4000 optimization runs in parallel for 5000 epochs (Fig 2C).

### Enhancement/suppression balance and decorrelation analysis

Our previous work revealed a relationship between decorrelation and how the enhancement/suppression balance in the V1 population evolves over time. To determine if this relationship is captured in the model, we calculated the enhancement/suppression index (ESI) and angle between SF vectors from model simulated responses (Fig 2D,E). Angles were chosen for this analysis because they provide a smooth measure of geometric separation over time, in contrast to correlations which can only be -1, 0, or 1 in a two-dimensional space. All estimates in this analysis were calculated across all 4000 model solutions; averages were calculated and standard deviations were reported as error measures (standard errors were deliberately avoided due to the large sample size making them uninformatively small).

To calculate the ESI, we first calculated the population-wide enhancement, *E(f*, *t)*, evoked by each SF by summing positive firing rates across the excitatory V1 population at each time lag. We calculated the sum of suppression, *S(f*, *t)*, in an identical manner by summing across negative firing rates. 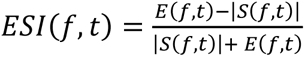 and is bounded between -1 and 1, where -1 denotes total suppression, 1 denotes total enhancement and 0 denotes a perfect balance between enhancement and suppression.

Angles between SF vectors was calculated as 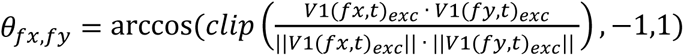, where *fx* and *fy* represent the population response evoked by two different SFs (x and y). Angles are directly related to correlations and inversely proportional: 0-degree angles correspond to perfectly correlated vectors, 90-degree angles correspond to perfectly uncorrelated/orthogonal vectors, and 180-degree angles correspond to perfectly anti-correlated vectors. Angles and ESI differences (dESI) were calculated between all SF pairs for comparison (Fig 2F).

### Model ablation experiments

Our previous work found strong evidence that both coarse-to-fine processing and intracortical inhibition are necessary for producing decorrelation. To test this hypothesis in the model, we performed three in-silico ablation experiments where parameters were selectively zeroed after fitting (Fig 3B-D). All estimates in these experiments were calculated in all 4000 model solutions. In the first experiment, we eliminated coarse-to-fine processing by setting the FTS parameter value to 0. In the second experiment, we eliminated intracortical inhibition by setting the INH_w_ parameter value to 0. In the final experiment, we eliminated both coarse-to-fine processing and intracortical inhibition by setting both parameters to 0.

### Fixed parameter optimization experiments

Three fixed-parameter optimization experiments were conducted in this study. Each of these was a separate optimization done with 1000 sets of initial conditions, 5000 training epochs, and the original parameter bounds. In the first experiment, FTS values were fixed to 0 while all other parameters were allowed to freely converge (Fig 3E). In the second experiment, INH_w_ values were fixed to 0, AMPG values were allowed to be negative (minimum of -4), and all other parameters were allowed to freely converge (Fig 3F). In the third experiment, AMPC was fixed to 0.16 while AMPW was fixed to 10 to produce a flat tuning curve (i.e., uniform peak firing rates across SF) while all other parameters were allowed to freely converge (Fig 5 A, B). We made statistical comparisons of the peak vector angles across all ablation and fixed parameter optimization conditions to quantify the effect of each manipulation (S1, supplementary information).

### Cross-parameter co-ordering permutation search

The values of different dLGN parameters can covary across circuit units in many ways, resulting in different permutations. For example, response latencies across afferents may increase or decrease with response amplitude gain. Because solutions can have different co-ordering permutations, averaging across solutions obscures their structure. Therefore, the model must be constrained to a single permutation to be structurally identifiable. To define this constraint, we applied a multi-step approach to determine which co-ordering permutation produced the lowest loss scores on average.

We first confirmed that the co-ordering permutations observed across the unconstrained model solutions were nonrandom. To do this, we quantified pairwise parameter covariance in each circuit by calculating the Spearman Autocorrelation Matrix (SAM) of each 7 x 3 dLGN parameter matrix. This makes a 7x7 matrix where each cell tells how positively or negatively the values of two parameters covary across afferents. An average across SAMs where the co-ordering permutations of the solutions are random would produce near 0 off-diagonals, as confirmed by a separate multi-start optimization in which random permutation matrices were applied to the dLGN parameter matrix during the forward pass (S2 Fig, panel i). The unconstrained solutions had an average SAM that differed systematically from random (S2 Fig, panel ii). The random permutations also produced higher loss scores, confirming that the optimizer converges onto permutations yielding lower loss (S2 Fig, panel iv).

Next, the permutations of the 120 lowest-loss unconstrained solutions were explicitly enforced in subsequent optimization procedure. A total of 100 optimization runs (5000 epochs) were done for each of the 120 permutations (S2 Fig, panel iii, same values as in Fig. 4G, L). The average loss of the best permutation was statistically similar to the next best 74 permutations (S2 Fig, panel v inset, vi). The permutation with the lowest average loss was selected (S2 Fig, panel v inset) and enforced in a final procedure with 4000 optimization runs and 10000 training epochs to yield an identifiable solution set, allowing functional interpretation of the circuit organization.

After applying the identifiability constraint, a local perturbation restart was applied to the solutions to determine whether the optimizer re-converged consistently onto a single global minimum. First, the parameter bounds were expanded beyond the original bounds (INH_w_: -10 - 10 Au; INH_d_: 0 – 100 ms; T: 0 – 200 ms; FTS: -200 – 200 ms; AMPM: -10 – 10 Au; AMPC: 0.01 – 1 c/d; AMPG: -10 – 10 Au; AMPW: 0.01 – 1 Au). Next, random noise from a uniform distribution (min = -1, max = 1) was added to the normalized parameters. The parameters were re-optimized for 1000 training epochs. The difference between parameter distributions before and after the perturbation restart was negligible (S2 Fig, panel v).

A separate parameter sweep analysis with expanded bounds was done to characterize the loss-landscape of the identifiable solution set, which confirmed that the minimum loss with respect to each parameter fell within the original bounds (S4 Fig, panel i, ii). The parameter sweep revealed that the AMPC and AMPW parameters were less constrained by the data; we reoptimized the model while progressively expanding the bounds for these parameters. Although the results were qualitatively similar, the original bounds still yielded the lowest average loss (S4 Fig, panel iii, vi).

### Fitting the model to V1 single units

Circuit instances were simultaneously fit to all 259 V1 single unit responses from our previous study (16) to infer the parameters of their dLGN inputs and compare them to in vivo dLGN estimates (Fig 6). The target V1 responses were averaged across orientation and phase conditions. For each of the 259 single units, 25 optimization runs were done for 20000 training epochs. When fitting with no identifiability constraints, the model overfit to the noise floor by converging onto AMPG values that made the overall amplitude of one or more dLGN unit responses near 0 (effectively silencing it). To address this, an identifiability constraint was implemented for all parameter classes such that param(dLGN_1_) ≤ param(dLGN_2_) ≤ param(dLGN_3_).

The T, FTS, and *σ* parameters from this optimization procedure were compared to the onset time, FTS, and response duration *in vivo* estimates respectively. The previous study determined response onset and offset times using a threshold equal to 2 standard deviations of the noise floor, where onset time was defined as the first threshold-crossing event, offset time was defined as the second threshold-crossing event and response duration was defined as the difference between offset and onset. The model predicted onset time was derived from T by subtracting 1.48*σ*, effectively estimating the threshold-crossing time by calculating its distance from the peak of the gaussian temporal envelope. The 1.48x scaling factor was derived analytically using the assumed signal-to-noise ratio (SnR) of 6 and noise floor of 2 STD from

Elsayed et al. (2025); 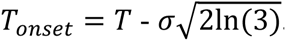. The response duration was similarly derived as 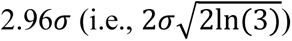, assuming a temporally symmetric response envelope.

### Statistical analysis

Comparisons of means were done via independent or related samples t-test (two-tailed) and comparisons of non-normal distributions were done with Kolmogorov-Smirnov test (two-tailed) using the SciPy package (version 1.13.1) in Python (version 3.9.23). Effect sizes were measured using a 3^rd^ party custom Cohen’s d Python function (68). All other Python code used is custom-written and available in the Data Availability Statement. Comparisons of Spearman autocorrelation matrices (SAMs) were done via Box’s M Test using the Statsmodels package (version 0.14.5) in Python. We did not perform a power analysis to predetermine sample sizes. We did not randomly assign animals to groups because it was not applicable to the experimental design of this study.

## Acknowledgments

We thank Dr. Rolf Skyberg for the electrophysiology data used in this study. We also thank Dr. Seiji Tanabe for discussion and consultation during our research. This work was supported by US National Institutes of Health (NIH) grants (EY026286 and EY020950 to J.C., EY032360 and R01DC018621 to C.D.M.), the National Science Foundation (IOS-1942480 to C.D.M.) and financial support from Jefferson Scholars Foundation to J.C.

## Data Availability Statement

All data and analysis code supporting the findings of this study are available on GitHub at https://github.com/Elsayedaa/synergistic_geniculocortical_circuit_model/tree/version_1.3

## Accession Numbers

Electrophysiological data used in above analysis will be posted to Figshare upon initial review.

## References

1. Olshausen BA, Field DJ. Emergence of simple-cell receptive field properties by learning a sparse code for natural images. Nature. 1996 Jun;381(6583):607–9. doi:10.1038/381607a0

2. Gutnisky DA, Dragoi V. Adaptive coding of visual information in neural populations. Nature. 2008 Mar;452(7184):220–4. doi:10.1038/nature06563

3. Benucci A, Ringach DL, Carandini M. Coding of stimulus sequences by population responses in visual cortex. Nat Neurosci. 2009 Oct;12(10):1317–24. doi:10.1038/nn.2398

4. Tanabe S. Population codes in the visual cortex. Neurosci Res. 2013 Jul;76(3):101–5. doi:10.1016/j.neures.2013.03.010 PubMed PMID: 23542219; PubMed Central PMCID: PMC3688279.

5. Shapiro JT, Gosselin EAR, Michaud NM, Crowder NA. Activating parvalbumin-expressing interneurons produces iceberg effects in mouse primary visual cortex neurons. Neuroscience Letters. 2022 Aug 24;786:136804. doi:10.1016/j.neulet.2022.136804

6. Zohary E. Population coding of visual stimuli by cortical neurons tuned to more than one dimension. Biol Cybern. 1992 Jan;66(3):265–72. doi:10.1007/BF00198480

7. Zohary E, Shadlen MN, Newsome WT. Correlated neuronal discharge rate and its implications for psychophysical performance. Nature. 1994 Jul 14;370(6485):140–3. doi:10.1038/370140a0 PubMed PMID: 8022482.

8. Stringer C, Pachitariu M, Steinmetz N, Carandini M, Harris KD. High-dimensional geometry of population responses in visual cortex. Nature. 2019 Jul;571(7765):361–5. doi:10.1038/s41586-019-1346-5

9. Kriegeskorte N, Diedrichsen J. Peeling the Onion of Brain Representations. Annu Rev Neurosci. 2019 Jul 8;42(1):407–32. doi:10.1146/annurev-neuro-080317-061906

10. Kriegeskorte N. Representational similarity analysis – connecting the branches of systems neuroscience. Front Sys Neurosci. 2008. doi:10.3389/neuro.06.004.2008

11. Kobak D, Brendel W, Constantinidis C, Feierstein CE, Kepecs A, Mainen ZF, et al. Demixed principal component analysis of neural population data. van Rossum MC, editor. eLife. 2016 Apr 12;5:e10989. doi:10.7554/eLife.10989

12. Cunningham JP, Yu BM. Dimensionality reduction for large-scale neural recordings. Nat Neurosci. 2014 Nov;17(11):1500–9. doi:10.1038/nn.3776

13. Vinje WE, Gallant JL. Sparse Coding and Decorrelation in Primary Visual Cortex During Natural Vision. Science. 2000 Feb 18;287(5456):1273–6. doi:10.1126/science.287.5456.1273

14. Matsumoto N, Okada M, Sugase-Miyamoto Y, Yamane S, Kawano K. Population Dynamics of Face-responsive Neurons in the Inferior Temporal Cortex. Cerebral Cortex. 2005 Aug 1;15(8):1103–12. doi:10.1093/cercor/bhh209

15. Roth ZN. Functional MRI Representational Similarity Analysis Reveals a Dissociation between Discriminative and Relative Location Information in the Human Visual System. Front Integr Neurosci. 2016 Mar 30;10. doi:10.3389/fnint.2016.00016

16. Elsayed A, Skyberg R, Cang J, Meliza CD. Synergistic Geniculate and Cortical Dynamics Facilitate a Decorrelated Spatial Frequency Code in the Early Visual System. J Neurosci. 2025 Nov 13;46(1). doi:10.1523/JNEUROSCI.1070-25.2025 PubMed PMID: 41233286.

17. McAdams CJ, Maunsell JHR. Effects of Attention on the Reliability of Individual Neurons in Monkey Visual Cortex. Neuron. 1999 Aug 1;23(4):765–73. doi:10.1016/S0896-6273(01)80034-9 PubMed PMID: 10482242.

18. Vreysen S, Zhang B, Chino YM, Arckens L, Van den Bergh G. Dynamics of spatial frequency tuning in mouse visual cortex. Journal of Neurophysiology. 2012 Jun;107(11):2937–49. doi:10.1152/jn.00022.2012

19. Allen EA, Freeman RD. Dynamic Spatial Processing Originates in Early Visual Pathways. J Neurosci. 2006 Nov 8;26(45):11763–74. doi:10.1523/JNEUROSCI.3297-06.2006 PubMed PMID: 17093097.

20. Nirody JA. Development of spatial coarse-to-fine processing in the visual pathway. J Comput Neurosci. 2014 Jun;36(3):401–14. doi:10.1007/s10827-013-0480-6 PubMed PMID: 24077933.

21. Bredfeldt CE, Ringach DL. Dynamics of Spatial Frequency Tuning in Macaque V1. J Neurosci. 2002 Mar 1;22(5):1976–84. doi:10.1523/JNEUROSCI.22-05-01976.2002 PubMed PMID: 11880528.

22. Schuurmans JP, Bennett MA, Petras K, Goffaux V. Backward masking reveals coarse-to-fine dynamics in human V1. Neuroimage. 2023 Jul 1;274:120139. doi:10.1016/j.neuroimage.2023.120139 PubMed PMID: 37137434.

23. Skyberg R, Tanabe S, Chen H, Cang J. Coarse-to-fine processing drives the efficient coding of natural scenes in mouse visual cortex. Cell Reports. 2022 Mar 29;38(13). doi:10.1016/j.celrep.2022.110606 PubMed PMID: 35354030.

24. Cai D, Deangelis GC, Freeman RD. Spatiotemporal Receptive Field Organization in the Lateral Geniculate Nucleus of Cats and Kittens. Journal of Neurophysiology. 1997 Aug;78(2):1045–61. doi:10.1152/jn.1997.78.2.1045

25. Ruksenas O, Bulatov A, Heggelund P. Dynamics of spatial resolution of single units in the lateral geniculate nucleus of cat during brief visual stimulation. J Neurophysiol. 2007 Feb;97(2):1445–56. doi:10.1152/jn.01338.2005 PubMed PMID: 16914606.

26. Parker PRL, Martins DM, Leonard ESP, Casey NM, Sharp SL, Abe ETT, et al. A dynamic sequence of visual processing initiated by gaze shifts. Nat Neurosci. 2023 Dec;26(12):2192–202. doi:10.1038/s41593-023-01481-7

27. Bellet J, Siegel M, Dehaene S, Jarraya B, Panagiotaropoulos T, Van Kerkoerle T. From Coarse to Rich: Successive Waves of Visual Perception in Prefrontal Cortex [Internet]. Neuroscience; 2026 [cited 2026 Jun 19]. Available from: http://biorxiv.org/lookup/doi/10.64898/2026.03.27.714202 doi:10.64898/2026.03.27.714202

28. Ringach DL, Bredfeldt CE, Shapley RM, Hawken MJ. Suppression of Neural Responses to Nonoptimal Stimuli Correlates With Tuning Selectivity in Macaque V1. Journal of Neurophysiology. 2002 Feb;87(2):1018–27. doi:10.1152/jn.00614.2001

29. Shapley R, Hawken M, Ringach DL. Dynamics of Orientation Selectivity in the Primary Visual Cortex and the Importance of Cortical Inhibition. Neuron. 2003 Jun 5;38(5):689–99. doi:10.1016/S0896-6273(03)00332-5 PubMed PMID: 12797955.

30. Levy M, Fournier J, Frégnac Y. The Role of Delayed Suppression in Slow and Fast Contrast Adaptation in V1 Simple Cells. J Neurosci. 2013 Apr 10;33(15):6388–400. doi:10.1523/JNEUROSCI.3609-12.2013 PubMed PMID: 23575837.

31. Olshausen BA, Field DJ. Emergence of simple-cell receptive field properties by learning a sparse code for natural images. Nature. 1996 Jun;381 (6583):607–9. doi:10.1038/381607a0

32. Simoncelli EP, Olshausen BA. Natural Image Statistics and Neural Representation. Annual Review of Neuroscience. 2001 Mar 1;24(Volume 24, 2001):1193–216. doi:10.1146/annurev.neuro.24.1.1193

33. Bell AJ, Sejnowski TJ. The “independent components” of natural scenes are edge filters. Vision Research. 1997 Dec;37(23):3327–38. doi:10.1016/S0042-6989(97)00121-1

34. Barlow HB, Földiák P. Adaptation and decorrelation in the cortex. In: The Computing Neuron. 1989. p. 54–72.

35. Ghodrati M, Khaligh-Razavi SM, Lehky SR. Towards building a more complex view of the lateral geniculate nucleus: Recent advances in understanding its role. Progress in Neurobiology. 2017 Sep 1;156:214–55. doi:10.1016/j.pneurobio.2017.06.002

36. Histed MH. Feedforward Inhibition Allows Input Summation to Vary in Recurrent Cortical Networks. eNeuro. 2018 Jan 1;5(1). doi:10.1523/ENEURO.0356-17.2018 PubMed PMID: 29682603.

37. Heggelund P, Hartveit E. Neurotransmitter receptors mediating excitatory input to cells in the cat lateral geniculate nucleus. I. Lagged cells. J Neurophysiol. 1990 Jun;63(6):1347–60. doi:10.1152/jn.1990.63.6.1347 PubMed PMID: 2162923.

38. Vigeland LE, Contreras D, Palmer LA. Synaptic Mechanisms of Temporal Diversity in the Lateral Geniculate Nucleus of the Thalamus. J Neurosci. 2013 Jan 30;33(5):1887–96. doi:10.1523/JNEUROSCI.4046-12.2013 PubMed PMID: 23365228.

39. Eichhorn J, Sinz F, Bethge M. Natural Image Coding in V1: How Much Use Is Orientation Selectivity? [Internet]. doi:10.1371/journal.pcbi.1000336

40. Failor SW, Carandini M, Harris KD. Visual experience orthogonalizes visual cortical stimulus responses via population code transformation. Cell Reports. 2025 Feb;44(2):115235. doi:10.1016/j.celrep.2025.115235

41. Chaudhuri R, Gerçek B, Pandey B, Peyrache A, Fiete I. The intrinsic attractor manifold and population dynamics of a canonical cognitive circuit across waking and sleep. Nat Neurosci. 2019 Sep;22(9):1512–20. doi:10.1038/s41593-019-0460-x

42. Stringer C, Michaelos M, Tsyboulski D, Lindo SE, Pachitariu M. High-precision coding in visual cortex. Cell. 2021 May 13;184(10):2767–2778.e15. doi:10.1016/j.cell.2021.03.042 PubMed PMID: 33857423.

43. Field DJ. Relations between the statistics of natural images and the response properties of cortical cells. J Opt Soc Am A. 1987 Dec 1;4(12):2379. doi:10.1364/JOSAA.4.002379

44. Field DJ. What Is the Goal of Sensory Coding? Neural Computation. 1994 Jul;6(4):559–601. doi:10.1162/neco.1994.6.4.559

45. Weliky M, Fiser J, Hunt RH, Wagner DN. Coding of Natural Scenes in Primary Visual Cortex. Neuron. 2003 Feb;37(4):703–18. doi:10.1016/S0896-6273(03)00022-9

46. Chalk M, Marre O, Tkačik G. Toward a unified theory of efficient, predictive, and sparse coding. Proc Natl Acad Sci USA. 2018 Jan 2;115(1):186–91. doi:10.1073/pnas.1711114115

47. Pouget A, Dayan P, Zemel R. Information processing with population codes. Nat Rev Neurosci. 2000 Nov;1(2):125–32. doi:10.1038/35039062

48. Mastronarde DN. Two classes of single-input X-cells in cat lateral geniculate nucleus. I. Receptive-field properties and classification of cells. Journal of Neurophysiology. 1987 Feb;57(2):357–80. doi:10.1152/jn.1987.57.2.357

49. Alonso JM, Usrey WM, Reid RC. Rules of Connectivity between Geniculate Cells and Simple Cells in Cat Primary Visual Cortex. J Neurosci. 2001 Jun 1;21(11):4002–15. doi:10.1523/JNEUROSCI.21-11-04002.2001 PubMed PMID: 11356887.

50. Yeh CI, Stoelzel CR, Alonso JM. Two Different Types of Y Cells in the Cat Lateral Geniculate Nucleus. Journal of Neurophysiology. 2003 Sep;90(3):1852–64. doi:10.1152/jn.00417.2003

51. Dong DW, Atick JJ. Temporal decorrelation: a theory of lagged and nonlagged responses in the lateral geniculate nucleus. Network: Computation in Neural Systems. 1995 Jan;6(2):159–78. doi:10.1088/0954-898X_6_2_003

52. Weng C, Yeh CI, Stoelzel CR, Alonso JM. Receptive Field Size and Response Latency Are Correlated Within the Cat Visual Thalamus. Journal of Neurophysiology. 2005 Jun;93(6):3537–47. doi:10.1152/jn.00847.2004

53. Lu J, Tucciarone J, Lin Y, Huang ZJ. Input-specific maturation of synaptic dynamics of parvalbumin interneurons in primary visual cortex. Proceedings of the National Academy of Sciences. 2014 Nov 25;111(47):16895–900. doi:10.1073/pnas.1400694111

54. Kukovska L, Wilmes KA, Homma NY, Clopath C, Poort J. Context-dependent activation of V1 parvalbumin interneurons enhances visual discrimination. PLOS Biology. 2025 Nov 25;23(11):e3003518. doi:10.1371/journal.pbio.3003518

55. Jadi MP, Sejnowski TJ. Regulating Cortical Oscillations in an Inhibition-Stabilized Network. Proceedings of the IEEE. 2014 May;102(5):830–42. doi:10.1109/JPROC.2014.2313113

56. Ozeki H, Finn IM, Schaffer ES, Miller KD, Ferster D. Inhibitory Stabilization of the Cortical Network Underlies Visual Surround Suppression. Neuron. 2009 May 28;62(4):578–92. doi:10.1016/j.neuron.2009.03.028 PubMed PMID: 19477158; PubMed Central PMCID: PMC2691725.

57. Mitchell JF, Sundberg KA, Reynolds JH. Spatial Attention Decorrelates Intrinsic Activity Fluctuations in Macaque Area V4. Neuron. 2009 Sep 24;63(6):879–88. doi:10.1016/j.neuron.2009.09.013 PubMed PMID: 19778515.

58. Cohen MR, Maunsell JHR. Attention improves performance primarily by reducing interneuronal correlations. Nat Neurosci. 2009 Dec;12(12):1594–600. doi:10.1038/nn.2439

59. Ruff DA, Cohen MR. Simultaneous multi-area recordings suggest that attention improves performance by reshaping stimulus representations. Nat Neurosci. 2019 Oct;22(10):1669–76. doi:10.1038/s41593-019-0477-1

60. Einevoll GT, Jurkus P, Heggelund P. Coarse-to-Fine Changes of Receptive Fields in Lateral Geniculate Nucleus Have a Transient and a Sustained Component That Depend on Distinct Mechanisms. PLOS ONE. 2011 Sep 9;6(9):e24523. doi:10.1371/journal.pone.0024523

61. Freund TF, Martin KA, Somogyi P, Whitteridge D. Innervation of cat visual areas 17 and 18 by physiologically identified X- and Y-type thalamic afferents. II. Identification of postsynaptic targets by GABA immunocytochemistry and Golgi impregnation. J Comp Neurol. 1985 Dec 8;242(2):275–91. doi:10.1002/cne.902420209 PubMed PMID: 2418072.

62. Tanaka K. Organization of geniculate inputs to visual cortical cells in the cat. Vision Research. 1985 Jan 1;Neural Basis of Visual Perception25(3):357–64. doi:10.1016/0042-6989(85)90060-4

63. Peters A, Payne BR. Numerical Relationships between Geniculocortical Afferents and Pyramidal Cell Modules in Cat Primary Visual Cortex. Cereb Cortex. 1993;3(1):69–78. doi:10.1093/cercor/3.1.69

64. Sedigh-Sarvestani M, Palmer LA, Contreras D. Thalamocortical synapses in the cat visual system in vivo are weak and unreliable. Huguenard J, Calabrese RL, editors. eLife. 2019 Apr 29;8:e41925. doi:10.7554/eLife.41925

65. Fröhlich F, Theis FJ, Hasenauer J. Uncertainty Analysis for Non-identifiable Dynamical Systems: Profile Likelihoods, Bootstrapping and More. In: Mendes P, Dada JO, Smallbone K, editors. Computational Methods in Systems Biology [Internet]. Cham: Springer International Publishing; 2014 [cited 2026 Aug 14]. p. 61–72. (Lecture Notes in Computer Science). Available from: https://link.springer.com/10.1007/978-3-319-12982-2_5 doi:10.1007/978-3-319-12982-2_5

66. Busse L, Wade AR, Carandini M. Representation of Concurrent Stimuli by Population Activity in Visual Cortex. Neuron. 2009 Dec 24;64(6):931–42. doi:10.1016/j.neuron.2009.11.004

67. Ruderman DL, Bialek W. Statistics of natural images: Scaling in the woods. Phys Rev Lett. 1994 Aug 8;73(6):814–7. doi:10.1103/PhysRevLett.73.814

68. skynaut. Answer to “How to calculate Cohen’s d in Python?” Stack Overflow [Internet]. Available from: https://stackoverflow.com/questions/21532471/how-to-calculate-cohens-d-in-python

