## Supplemental Figure S1 for "A geniculocortical circuit model predicts functional organization underlying efficient spatial frequency coding in the mouse visual system"

**S1. Decorrelation changes under different model ablation and fixed optimization conditions**

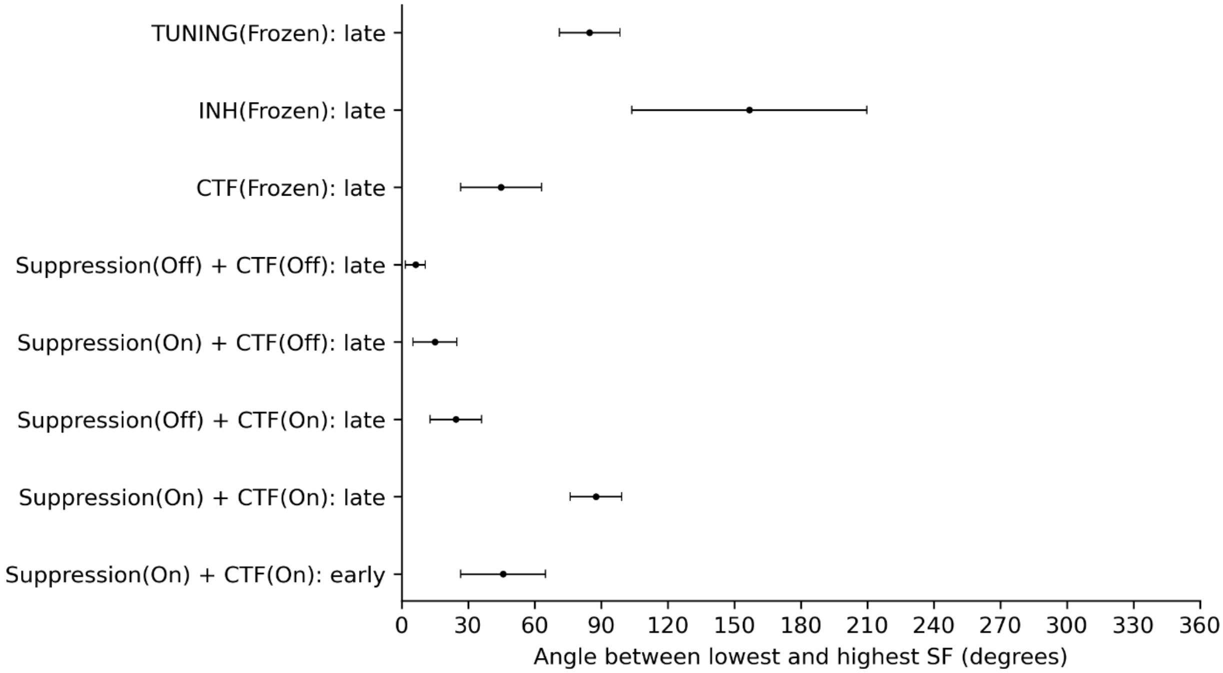


Figure A: **Angle between lowest and highest SF vector in different ablation or fixed optimization conditions:** Top 6 rows: Late phase (peak angles) of different ablation and fixed optimization conditions. Bottom 2 rows: Vector angles in the baseline condition (Suppression(On) + CTF(On)) during the late phase (peak angle) and early phase (peak dot product).

| condition | Compared to | T_statistic | pval | df | cohensd |
|---|---|---|---|---|---|
| Suppression(Off) + CTF(On) | Early control | 60.281613 | 0.000000e+00 | 7998.0 | 1.347938 |
| Suppression(Off) + CTF(On) | Late control | 243.326419 | 0.000000e+00 | 7998.0 | 5.440944 |
| Suppression(On) + CTF(Off) | Early control | 90.145136 | 0.000000e+00 | 7998.0 | 2.015707 |
| Suppression(On) + CTF(Off) | Late control | 300.374786 | 0.000000e+00 | 7998.0 | 6.716584 |
| Suppression(Off) + CTF(Off) | Early control | 127.193392 | 0.000000e+00 | 7998.0 | 2.844131 |
| Suppression(Off) + CTF(Off) | Late control | 413.194686 | 0.000000e+00 | 7998.0 | 9.239314 |
| CTF(Frozen) | Early control | 1.231160 | 2.183209e-01 | 4998.0 | 0.043528 |
| CTF(Frozen) | Late control | 91.334039 | 0.000000e+00 | 4998.0 | 3.229146 |
| INH(Frozen) | Early control | -107.500137 | 0.000000e+00 | 4988.0 | -3.816029 |
| INH(Frozen) | Late control | -75.760272 | 0.000000e+00 | 4988.0 | -2.689331 |
| TUNING(Frozen) | Early control | -60.896305 | 0.000000e+00 | 4998.0 | -2.153010 |
| TUNING(Frozen) | Late control | 6.574434 | 5.382288e-11 | 4998.0 | 0.232441 |

Table A: **Statistical comparisons for the conditions in Figure A:** All experimental conditions are compared to the early phase and late phase controls. Lowest effect size (highest similarity) is between CTF(Frozen) vs. early control and TUNING(Frozen) vs. late control.
