## Supplemental Figure S2 for "A geniculocortical circuit model predicts functional organization underlying efficient spatial frequency coding in the mouse visual system"

### S2. Cross-parameter co-ordering permutation search

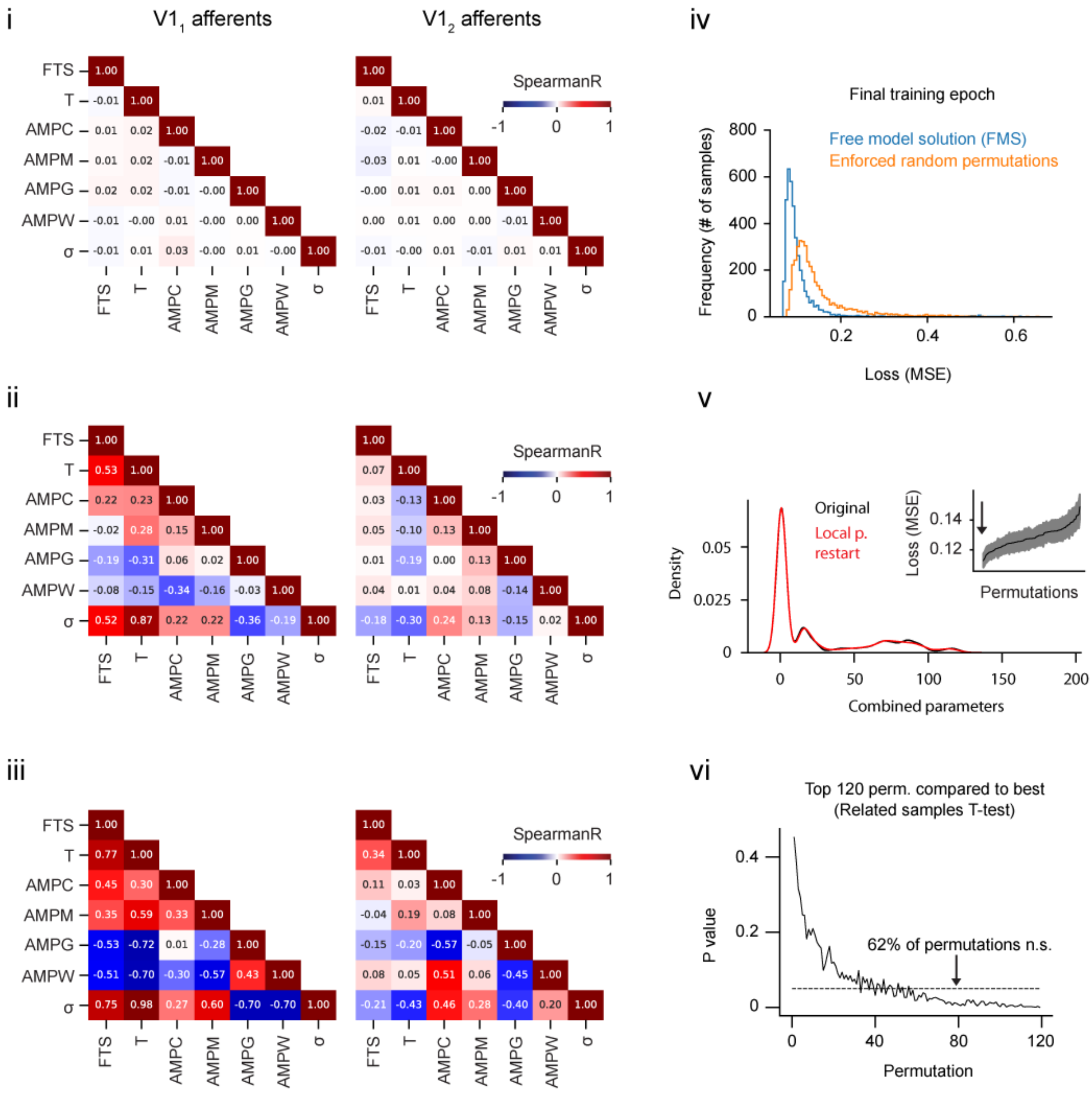


Figure B: **Permutation search analysis:** (i) Average (across optimization runs) Spearman autocorrelation matrix (SAM) for model optimized with random permutations. (ii) Average SAM for the original optimization condition with no permutation/identifiability constraint. Co-ordering of parameters across afferents is significantly different from random (Box's M, $X^2(28) = 803.62$, $p < 0.001$). (iii) Average SAM for the best 120 permutations (100 samples per permutation), identical to Fig. 4G, L but with annotated cells. (iv) Comparison of loss distributions from random permutation vs. unconstrained optimization. Average loss for the unconstrained optimization is significantly lower (Independent T-test, statistic=-30.4, p < 0.001, df=7998, Cohen's d = -0.68). (v) Kernel density estimate of all model parameter values before (black) and after local perturbation restart (red). The difference is negligible (Kolmogorov-Smirnoff test, statistic=0.016, p < 0.0001, df =167999, Cohen's d < 0.002). Inset: Average loss across the top 120 permutations identified in the permutation search (See methods: Cross-parameter co-ordering permutation search). (vi) The 120 permutations with the lowest loss compared to the best permutation (lowest overall average loss). The average loss of the best permutation was not significantly different from the next best 74 permutations (Related samples T-test, p>0.05, df = 99).
