## Supplemental Figure S3 for "A geniculocortical circuit model predicts functional organization underlying efficient spatial frequency coding in the mouse visual system"

### S3. Spearman autocorrelation matrices with flat amplitude tuning curves

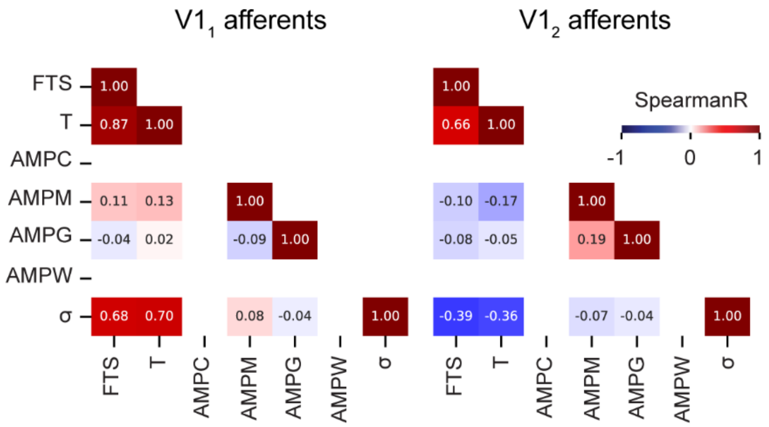


Figure C: **Spearman autocorrelation matrices with flat amplitude tuning curves:** AMPC and AMPW parameters were set to values that produce a flat amplitude tuning curve. All other parameters are left to freely converge. The figure shows the SAM from the flat tuning optimization.
