## Supplemental Figure S4 for "A geniculocortical circuit model predicts functional organization underlying efficient spatial frequency coding in the mouse visual system"

**S4. Parameter sweep loss analysis**

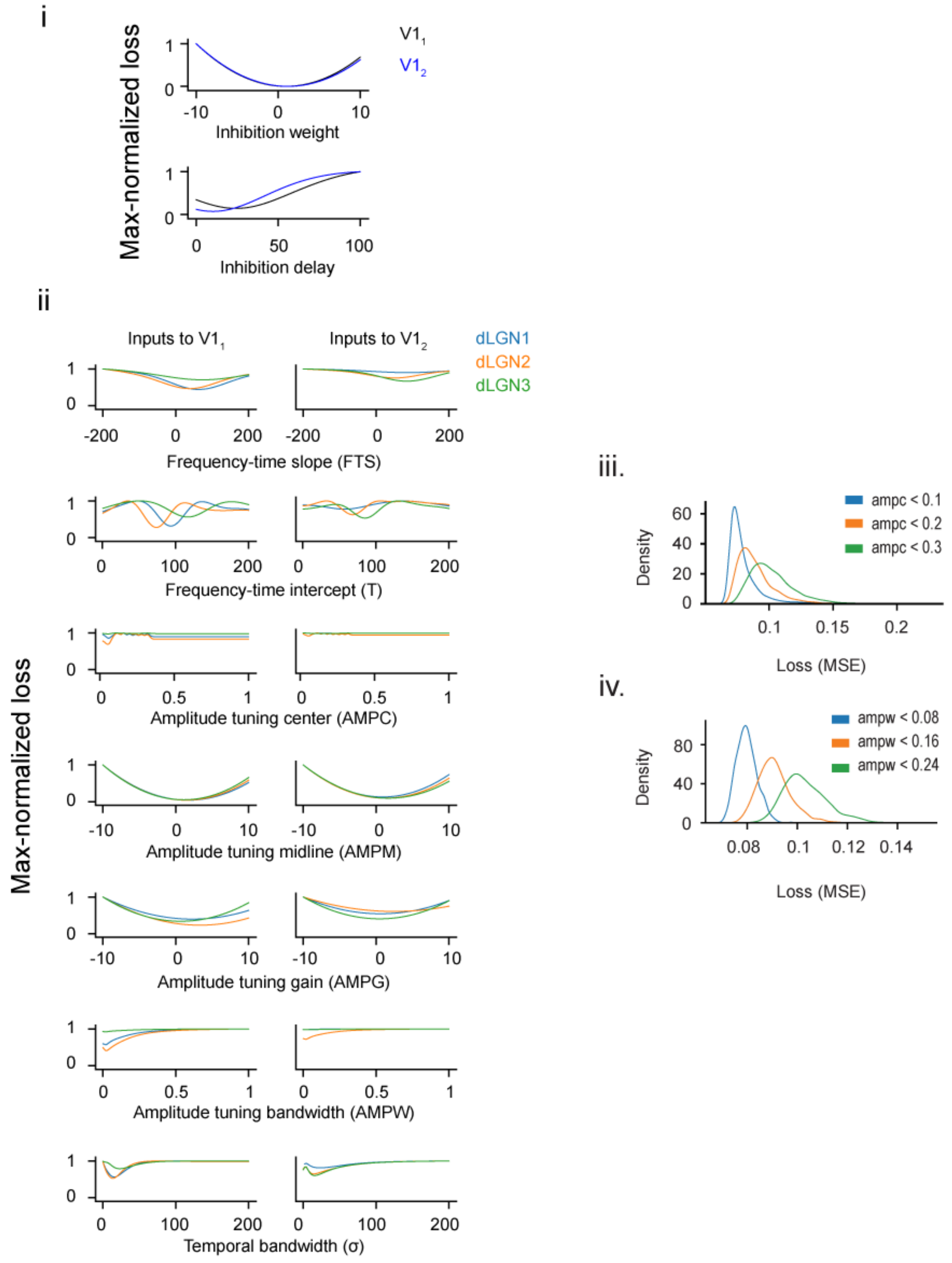


Figure D: **Characterization of the loss landscape with respect to model parameters:** (i) Max-normalized loss with respect to the V1 parameters. (ii) Max-normalized loss with respect to the dLGN parameters. (iii) Loss distributions across optimization runs for each of the AMPC parameter expanded bound conditions (Independent T-test, statistic < -36.4, df =

1998; $p < 0.0001$, Cohen's $d < 1.62$ for all comparisons). (iv) Same as A. but for the AMPW parameter (statistic < -36.4, df = 1998; $p < 0.0001$, Cohen's $d < 1.63$ for all comparisons).
